# Neuron-level Prediction and Noise can Implement Flexible Reward-Seeking Behavior

**DOI:** 10.1101/2024.05.22.595306

**Authors:** Chenguang Li, Jonah Brenner, Adam Boesky, Sharad Ramanathan, Gabriel Kreiman

## Abstract

We show that neural networks can implement reward-seeking behavior using only local predictive updates and internal noise. These networks are capable of autonomous interaction with an environment and can switch between explore and exploit behavior, which we show is governed by attractor dynamics. Networks can adapt to changes in their architectures, environments, or motor interfaces without any external control signals. When networks have a choice between different tasks, they can form preferences that depend on patterns of noise and initialization, and we show that these preferences can be biased by network architectures or by changing learning rates. Our algorithm presents a flexible, biologically plausible way of interacting with environments without requiring an explicit environmental reward function, allowing for behavior that is both highly adaptable and autonomous. Code is available at https://github.com/ccli3896/PaN.

## 1 Introduction

There is a growing consensus that nervous systems are emergent [44, 37], meaning that behaviors and computations emerge from the interactions of many small components (neurons) rather than the independent actions of larger modules. Neuron-level prediction is a biologically plausible theory of how neural networks form useful sensory representations [33, 15], and together with the emergent view of the brain, it is likely that neurons update their activities and connection strengths with a small and consistent set of rules that include a predictive component [13, 19, 22, 28].

However, predictive agents are susceptible to the “dark room problem,” where agents minimize predictive errors by either reducing their activity to zero or staying in places where nothing happens [42]. Predictive agents that explore and act in environments usually need additional components to work, such as separate action selection modules [13, 38] or curiosity drives [18].

At the same time, biophysical studies have long established the presence of noise in nervous systems. Sources of noise include ion channel fluctuations, synaptic failure, or inconsistent vesicular release at synapses [11, 12, 25]. The role of noise in the brain is still not agreed upon, especially considering that some forms of it—such as spontaneous neural activity—can be energetically expensive to maintain [26]. Common perspectives are that nervous systems must somehow compensate for noise, average over it, or use noise as a means of regularization [12, 27].

Here we propose a new role for internal noise. We show that noise itself can overcome the dark room problem while adhering to the predictive and emergent view of behavior. We combine noise with predictive coding models [32, 39, 34, 41] to form our algorithm PaN (Prediction and Noise), and find that it can interact with environments in interesting ways. Our main contributions are to show that:

- Local prediction and noise alone can implement reward-seeking behavior.
- Networks switch between exploration and exploitation phases without prompting. These phases can be explained by attractor dynamics.
- Networks adapt to both internal and external changes. Internal change refers to the addition or removal of neurons; external change refers to environments with dynamic reward structures or motor interfaces.
- Networks vary in their goal preferences, which can be biased in biologically relevant ways: initialization, experience, architecture, and modulated learning rates.

PaN behavior is emergent rather than governed by optimization of an environmental reward. It therefore differs from the classical reinforcement learning framework, and so we place PaN under the novel framework of *self-supervised behavior* (SSB), motivated and defined in Appendix A.

## 2 Related work

### 2.1 Prediction as action

Earlier work in active inference [20, 17] has suggested that actions can be seen as a way to minimize predictive error, illustrated in Figure 1. Parr and Pezzulo [36] state that “active inference is a normative framework to characterize Bayes-optimal behavior and cognition in living organisms,” suggesting that “living organisms follow a unique imperative: minimizing the surprise of their sensory observations.”

**Figure 1.**
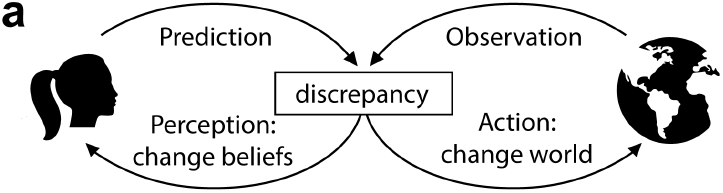
**a**. Active inference, adapted from Figure 2.3 of Parr and Pezzulo [36]. When an organism makes predictions about its environment, it may experience discrepancies with its observations. To reduce discrepancies, the organism can either change its beliefs about the environment or change the environment to match its predictions.

However, most active inference models require components with separate roles [31, 36, 17, 14] and thus do not operate on emergent principles. Active inference models are computationally costly and critics have remarked that they can be difficult to build or understand [6, 16, 4]. Although these models have been successful in certain environments [23, 31, 46], one must still specify learned target distributions, and scaling them up is an ongoing problem [45].

We use the idea from active inference that actions are a way to minimize predictive errors. But we do not aim to explain or generate Bayes-optimal behavior—nor do we aim to produce behavior that is optimal with respect to any predefined utility functions. Moreover, the continuous injection of internal noise in our model both *prevents* optimality in static environments and *is necessary* for most of our listed contributions (Section 4). Authors of active inference and the related free energy principle literature, on the other hand, assume that one can average over internal noise to arrive at long-term behavior [16].

### 2.2 Relevant models

We build on the existing body of work in predictive coding (PC), especially Monte Carlo Predictive Coding (MCPC) [34] and Incremental Predictive Coding (iPC) [39], both reviewed briefly here. While previous work has focused on PC’s role in perception or the learning of fixed targets, we demonstrate novel capabilities of these algorithms in actively behaving contexts.

#### Base Predictive Coding algorithm

In PC as defined by Bogacz and colleagues [32, 41], networks minimize a predictive energy (Equation 1) [32] through activity and weight updates. To learn an input-output mapping, a PC network’s first and last layer’s activities are fixed to target values; then, network activities are updated to minimize prediction error. After activity updates are run to convergence, weights are updated for a single step to minimize the same prediction error. All updates use local information, and activity and weight updates alternate until the network has converged. Song et al. (2024) [41] show that this procedure may enable more efficient and biologically plausible learning than backpropagation, where loss-minimizing updates are restricted to weights.

#### Monte Carlo Predictive Coding

MCPC adds noise to the activity updates from PC [34]. Given images from the MNIST dataset, MCPC can infer and sample posterior distributions of latent variables. The algorithm is robust to a range of noise settings and can account for experimental findings; for example, it explains a reduction in neural variability after stimulus presentation, which has been observed across a variety of conditions in animals [7].

#### Activity and weight coupling

We also couple our activity and weight updates as in Incremental predictive coding (iPC) [39]. iPC modifies PC by updating activities and weights in alternation, so activities are no longer updated to convergence. Coupling eliminates the need in PC for an external control signal to switch the network from activity updates to the weight update step. iPC is faster than PC and has convergence guarantees, whereas convergence in PC can be unpredictable [39].

Activity and weight coupling has been studied in other contexts, such as recurrent neural network (RNN) training. While our models do not have recurrent weights, an effective recurrence can be introduced with environmental feedback. RNNs are normally trained to produce desired activity dynamics with frozen weights, but alternative training schemes using coupled activity and weight updates can impart computational benefits [30, 29, 21, 2]—in one example, Clark and Abbott (2024) [8] describe how such coupling can serve as a mechanism for working memory.

## 3 Methods

### Algorithm 1

Prediction and Noise (PaN).

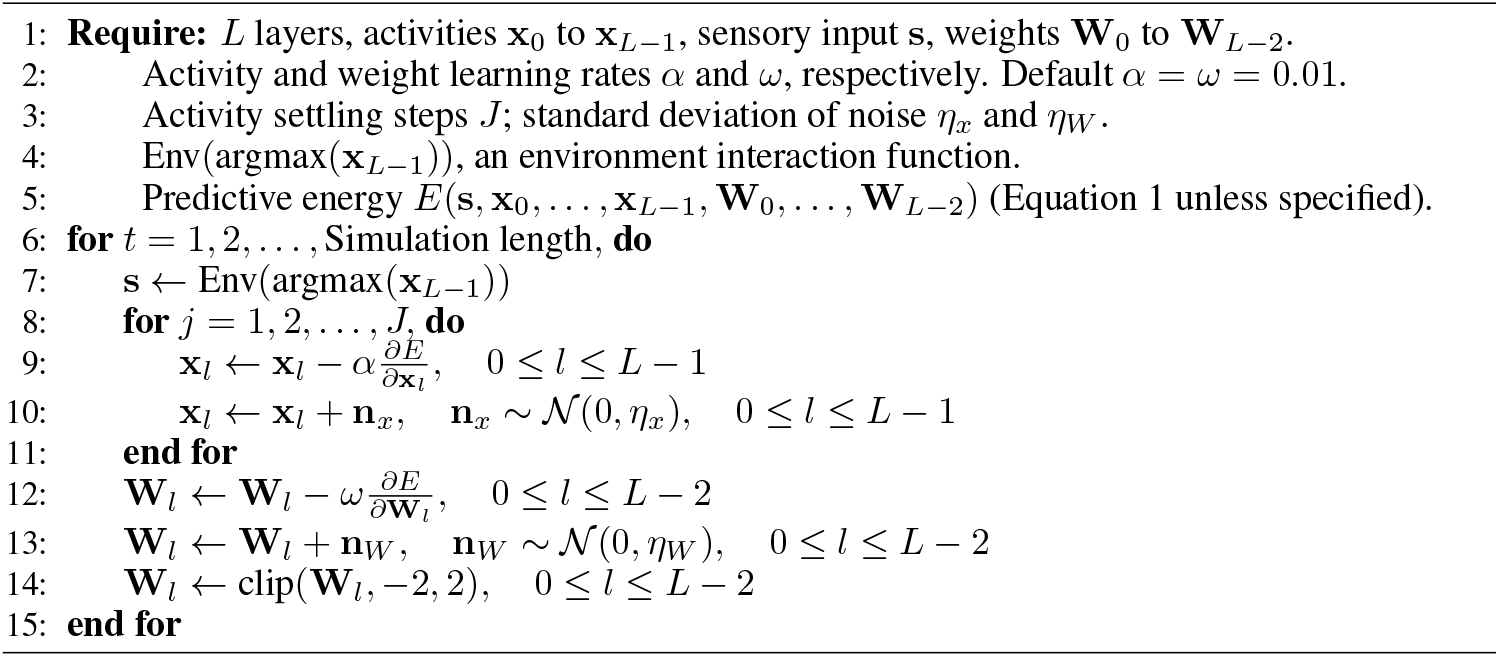

Figure 2a shows an example PaN network with an argmax-based environment function and fixed action-to-signal mappings. For a network of L layers, let the neuron activities for layer *l* be denoted **x**_*l*_ and the weights connecting layer *l* to *l* + 1 be **W**_*l*_. Let **s** be a scalar or vector of sensory feedback. We define a predictive energy for each timestep *t*:

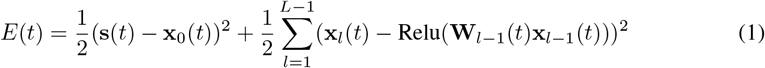

Relu is the rectified linear activation function. The sensory feedback **s**(*t*) is determined by some environment function Env(**x**_*L*−1_(*t* − 1)) = **s**(*t*); we specify this environment function in each experiment. Networks are updated using Algorithm 1, illustrated in Figure 2b. Activities and weights are updated in alternation to minimize the predictive energy (Equation 1). Every update is accompanied by the addition of white noise, independent at every timestep and normally distributed with a standard deviation of *η*_*x*_ for activities and *η*_*W*_ for weights. All experiments in the paper were run on Intel Cascade Lake or Sapphire Lake CPUs.

**Figure 2.**
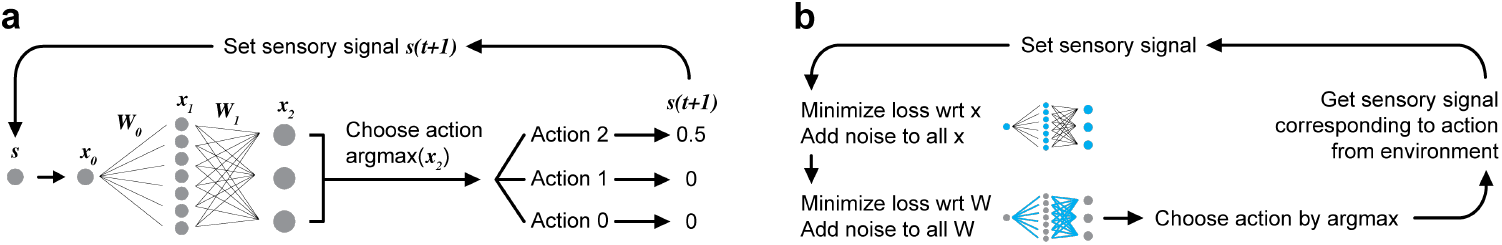
**a**. Flow of information in a Prediction and Noise (PaN) network, using an example set of possible action-to-signal mappings. **s** is an input, **x**_*l*_ denotes the activity vector for each layer *l*, and **W**_*l*_ denotes the weight matrix connecting layer *l* to *l* + 1. **b**. Update loop (see Algorithm 1).

## 4 Results

### 4.1 Two-neuron networks seek reward and switch between exploration and exploitation

We start with two neurons connected by a weight, Figure 3a, with no nonlinearity in the energy for ease of analysis:

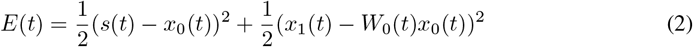

PaN networks are run for 500k timesteps per trial in the closed-loop setup in Figure 3a. Without noise, *η*_*x*_ = *η*_*W*_ = 0, networks randomly fixate on one action (Figure 3b). With noise, networks learn to prefer Action 1, which is associated with nonzero signal and in this context looks like reward-seeking behavior (Figure 3c-d). Looking more closely at an example noise setting *η*_*x*_ = 0.042, *η*_*W*_ = 0.0013 in Figure 3e-h, we see that action choices (Figure 3e) alternate between high entropy exploratory states and low entropy exploitative states. In contrast, trained ϵ-greedy agents have uniform patterns of exploration (Figure 3f,h). Appendix B shows that the separation of high and low entropy states is not strongly dependent on the window size used. Activity and weight noise scales impact the amount of reward collected (Figure 3i-j) with a maximum at *η*_*x*_ = 0.0075, *η*_*W*_ = 0 (see Appendix C Table 1) and different qualitative behaviors for different noise settings. See Appendix C for further examples.

**Figure 3.**
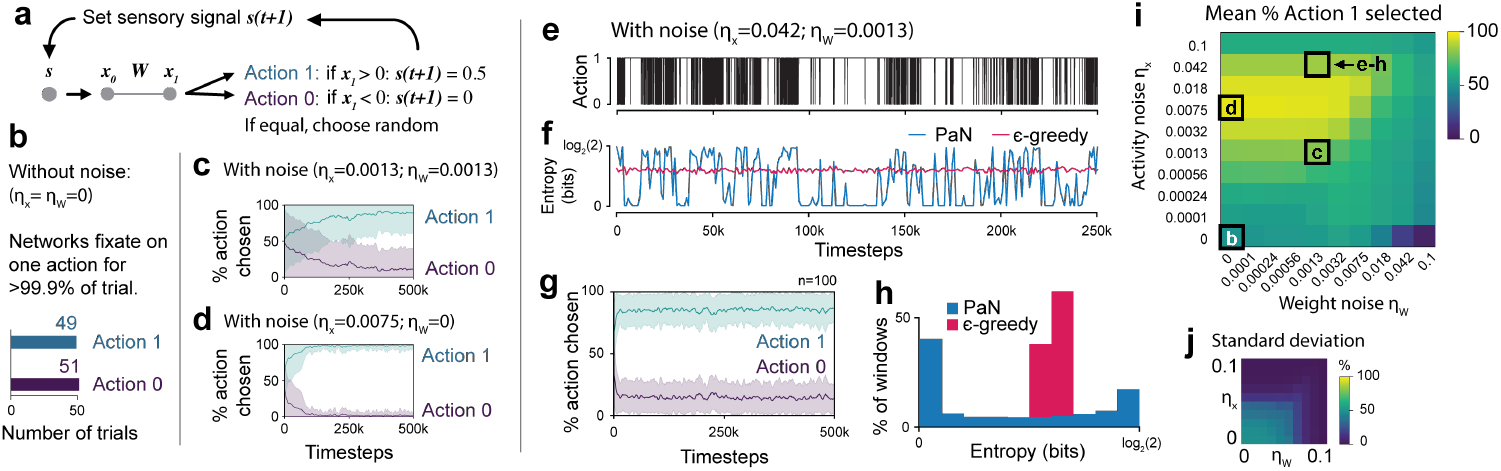
**a**. A two-neuron network that can take two actions. Updates follow Algorithm 1 without Relu nonlinearity for ease of analysis; J=1. **b**.. Without noise, networks randomly fixate on one action. **c-d**. With noise, networks choose the rewarding action at a rate dependent on activity and weight noise values. Mean plotted over 100 seeds. Shaded regions denote standard deviation. **e**. Sample actions chosen over 250k timesteps for *η*_*x*_ = 0.042, *η*_*W*_ = 0.0013 where *η* values define the width of noise distributions. **f**. Entropy of rolling 1000-timestep windows for PaN as well as an ϵ-greedy algorithm with ϵ set to 0.3 for matched reward 85%. **g**. Same as (c-d) but for *η*_*x*_ = 0.042, *η*_*W*_ = 0.0013. **h**. Histogram of entropy values over 100 seeds for 500k timesteps each. In this noise setting, PaN is bimodal with peaks corresponding to exploration (entropy close to log_2_(2) = 1) and exploitation (entropy close to 0). An ϵ-greedy agent, in contrast, maintains a consistently random exploration strategy. Appendix B shows that bimodality is not strongly dependent on window size. **i**. Different noise scales were tested for 100 seeds, 500k timesteps each. The mean percentage of time networks selected Action 1, the rewarding action, with standard deviations in **j**. See Appendix C Table 1 for values. Upper bound for compute for this figure and Figure 4 was 500 CPU hours, 55 GB for storage.

**Table 1:** Values for Figure 3i-j. **a**. Mean and **b**. standard deviation of proportion of actions taken that were rewarding for an action space with 2 actions corresponding to rewards [0, 0.5]. For two-neuron system in Figure 3a. Note that here, random actions would correspond to a mean of 0.5. The maximum means in (a) are bolded (.957).

| (a) |  |  |  |  |  |  |  |  |  |  |
| --- | --- | --- | --- | --- | --- | --- | --- | --- | --- | --- |
| $\eta_x \backslash \eta_W$ | 0 | .0001 | .00024 | .00056 | .0013 | .0032 | .0075 | .018 | .042 | .1 |
| 0 | .510 | .514 | .529 | .530 | .538 | .572 | .540 | .428 | .186 | .059 |
| .0001 | .556 | .554 | .552 | .552 | .576 | .623 | .615 | .582 | .555 | .457 |
| .00024 | .608 | .610 | .604 | .596 | .600 | .632 | .620 | .585 | .560 | .478 |
| .00056 | .718 | .721 | .723 | .716 | .669 | .657 | .633 | .590 | .565 | .496 |
| .0013 | .807 | .809 | .811 | .816 | .809 | .725 | .663 | .601 | .573 | .512 |
| .0032 | .915 | .915 | .915 | .915 | .911 | .877 | .733 | .628 | .588 | .530 |
| .0075 | **.957** | **.957** | **.957** | **.957** | .955 | .943 | .851 | .684 | .614 | .552 |
| .018 | .947 | .947 | .947 | .947 | .947 | .942 | .914 | .767 | .651 | .579 |
| .042 | .851 | .851 | .851 | .851 | .851 | .849 | .841 | .778 | .655 | .571 |
| .1 | .615 | .615 | .615 | .615 | .615 | .615 | .613 | .605 | .576 | .537 |

| (b) |  |  |  |  |  |  |  |  |  |  |
| --- | --- | --- | --- | --- | --- | --- | --- | --- | --- | --- |
| $\eta_x \backslash \eta_W$ | 0 | .0001 | .00024 | .00056 | .0013 | .0032 | .0075 | .018 | .042 | .1 |
| 0 | .500 | .495 | .488 | .475 | .434 | .379 | .217 | .099 | .036 | .013 |
| .0001 | .483 | .483 | .471 | .449 | .410 | .349 | .205 | .090 | .043 | .017 |
| .00024 | .455 | .453 | .447 | .429 | .398 | .345 | .204 | .090 | .043 | .017 |
| .00056 | .384 | .380 | .374 | .365 | .360 | .332 | .200 | .089 | .043 | .017 |
| .0013 | .329 | .326 | .321 | .306 | .275 | .281 | .191 | .088 | .042 | .017 |
| .0032 | .146 | .146 | .146 | .146 | .149 | .142 | .163 | .084 | .041 | .017 |
| .0075 | .048 | .048 | .048 | .048 | .051 | .055 | .086 | .075 | .038 | .016 |
| .018 | .022 | .022 | .022 | .022 | .023 | .024 | .032 | .055 | .034 | .015 |
| .042 | .018 | .018 | .018 | .018 | .018 | .019 | .020 | .029 | .026 | .013 |
| .1 | .018 | .018 | .018 | .018 | .018 | .018 | .018 | .018 | .016 | .012 |

#### 4.1.1 Attractor dynamics explain two-neuron behavior

For intuition as to why PaN selects the rewarding action, consider the case where Action 0 is chosen. Then *s*(*t* + 1) = 0, and using labels from Figure 3a, *x*_0_ and *x*_1_ will both quickly converge to 0. When *x*_1_ is close to 0, the next action is mostly determined by noise, leading to high prediction error. But when Action 1 is chosen, *s*(*t* + 1) = 0.5, and *x*_1_ will instead converge to 0.5*W*. As long as *W* is not close to 0, the network is likely to fixate on a single action, leading to low prediction error.

We can explain the explore/exploit behavior of the PaN network by studying the underlying dynamical system. When *s* in Equation 2 is fixed to either 0 or 0.5, we can derive an associated attractor in the parameter space, (*x*_0_, *x*_1_, *W*). Figure 4a shows the line attractor associated with no reward and Figure 4b the attractor with reward. Small plots in Figure 4a-b are the lines that networks move toward, and colored plots show how networks move, from yellow to purple.

**Figure 4.**
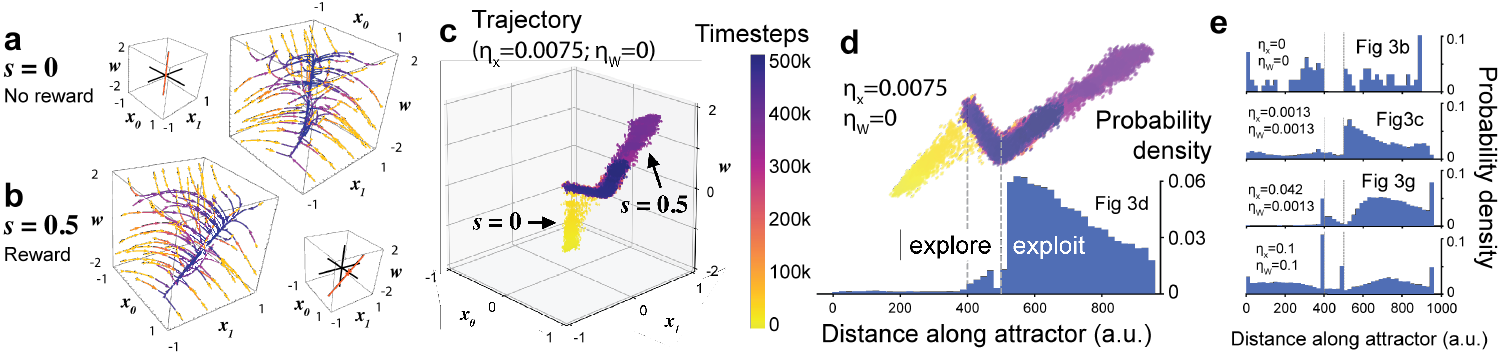
**a-b**. Attractors derived in Appendix D for the two-neuron system in Figure 3a. Small 3D plots are the line attractors for each case and the larger colored plots show how networks move toward attractors, from yellow to purple over time. For *s* = 0, the attractor is along the *W* -axis (a). For *s* = 0.5, the attractor is along *x*_1_ = *W x*_0_ (b). **c**. A single trajectory in parameter space (*x*_0_, *x*_1_, *W*) with movement along and between attractors. **d**. Probability density of 100 trajectories along attractor branches in arbitrary units (a.u.), with reference locations on the attractor marked in gray dashed lines. In this noise setting, networks strongly prefer the exploit branch. We only include timesteps 200k-500k to account for early transients in (d-e). **e**. Probability densities of other noise settings, 100 trials each. Without noise (top), networks stay along the part of the attractor where they began. With high noise (bottom), where networks distribute themselves widely and do not seek reward. Gray dashed lines correspond to the same locations as in (d).

A parameter space trajectory in Figure 4c for the noise setting in Figure 3d shows that the network moves along and between the attractors. Attractor stabilities are determined by analyzing the Jacobian of the stochastic differential equation approximation of the network, which we explain and empirically validate in Appendix D. The three segments of the trajectory correspond to *s* = 0 (Figure 4a), which is unstable in the presence of noise; *s* = 0.5 (Figure 4b), which is stable exploitation behavior; and movement between these two states, which is exploration behavior also driven by noise. We refer to the last segment as the attractor “bridge” for convenience. Video 1 shows the correspondence of trajectory location and explore/exploit behavior for the two-neuron system.

Figure 4d shows the probability density of 100 trajectories like the one in Figure 4c, plotting the densities along the closest point on the attractors and their bridge. In Figure 4d, the network is inclined toward exploitation. Changing probability densities in Figure 4e show that noise affects the tendency of a network to explore or exploit. Without noise, networks stay where they began and do not prefer one action over another (Figure 4e, top). At intermediate levels of noise (Figure 4d, e second and third plots), networks occupy the attractor corresponding to exploitation with higher probability. With very high noise (Figure 4e, bottom) networks distribute themselves between both attractors and do not show consistent reward-seeking. See Appendix D for details.

### 4.2 Larger PaN networks learn to chose maximally rewarding actions

Previous results show that in the presence of noise, a two-neuron system chooses a rewarding action more often than a zero-reward action. We next test a larger network with a 30-neuron hidden layer that can choose between six actions with linearly spaced rewards (Figure 5a). Networks learn to choose the maximal action over time (Figure 5b, with c as an example trace). As in Figure 3h, Figure 5e shows greater variation in entropy distributions as compared to a matched-reward ϵ-greedy agent.

**Figure 5.**
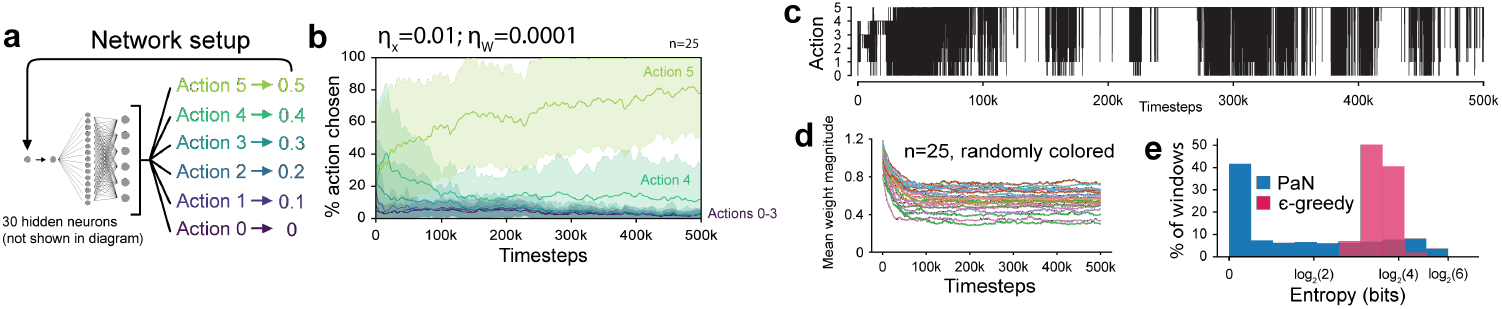
PaN networks learn to maximize sensory signals. **a**. PaN networks with linearly spaced action choices **b**. learn to optimize signal, not just choose nonzero values. Mean and standard deviation plotted over 25 seeds. **c**. Example trace for (a-b). **d**. Signal optimization can be explained by weight decay, automatically implemented by PaN loss (see text for discussion). Each line is the mean weight magnitude 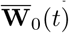 over the course of the trial. 25 random seeds. **e**. Entropy distributions for PaN networks in (a-d), 25 trials, with distributions for 25 trials of matched-reward ϵ-greedy agent activity. Experiments required upper bound of 6.25 CPU hours, 10 GB.

To provide intuition for why PaN fixates on actions that maximize signals rather than any nonzero action, note that the predictive energy increases through both fluctuating inputs and internal noise. The contribution through input fluctuation can be eliminated by making a set of weights large enough to fixate on one action, and the contribution through internal noise can be minimized by shrinking all weights. Because larger inputs allow for smaller weights to achieve fixation, networks prefer actions that lead to the largest input signals. Consistent with this intuition, we observe that weight decay is a consequence of minimizing the PaN loss in the presence of noise (Figure 5d).

### 4.3 PaN adapts to both internal and environmental changes

#### 4.3.1 PaN automatically adjusts to architectural changes

PaN can adapt itself to changing hidden layer sizes in the bandit task from Figure 5a. In Appendix E, Figure 14a-b, we first show that networks with 20 or 40 hidden neurons can still learn the optimization task. Then in Figure 14c, we run networks with 40 hidden neurons and randomly cut 20 neurons at 500k timesteps, showing that networks adapt and then continue to select the highest-reward action most of the time. Similar adaptation is shown in Figure 14d, where networks start with 20 hidden neurons and are then doubled to 40 neurons at 500k timesteps.

#### 4.3.2 PaN adapts to external changes

##### Direct action-to-reward mappings

We now test PaN’s performance as a continual learning agent using dynamic environments [1]. Appendix Figure 14e again begins with the 6-action setup in Figure 5a, now with 30 neurons in the hidden layer. At 250k timesteps, the reward associations are flipped—the action that was previously most rewarding is now the least rewarding, and vice versa. PaN autonomously learns to exploit the new best action over the following 500k timesteps.

##### Open-field search tasks

For a more complex environment, we design a task where actions correspond to movements in a reward landscape (6a). Instead of fixed rewards, we set the change in reward between an agent’s previous location and its new location as the input *s*. This one-timestep difference mimics an immediate “adaptation” at the sensory level to the previous reward value. When agents are placed in the landscape in Figure 6b, they spend more time at locations with greater reward values (sample track in Figure 6c; occupation density in Figure 6d). Agents do not fixate on one local maximum but usually explore multiple peaks during a single trial (Figure 6e). In fact, most agents initialized at random locations spent at least 10% of the trial in at least 3 different quadrants of the landscape, and many agents spent time in all 4.

**Figure 6.**
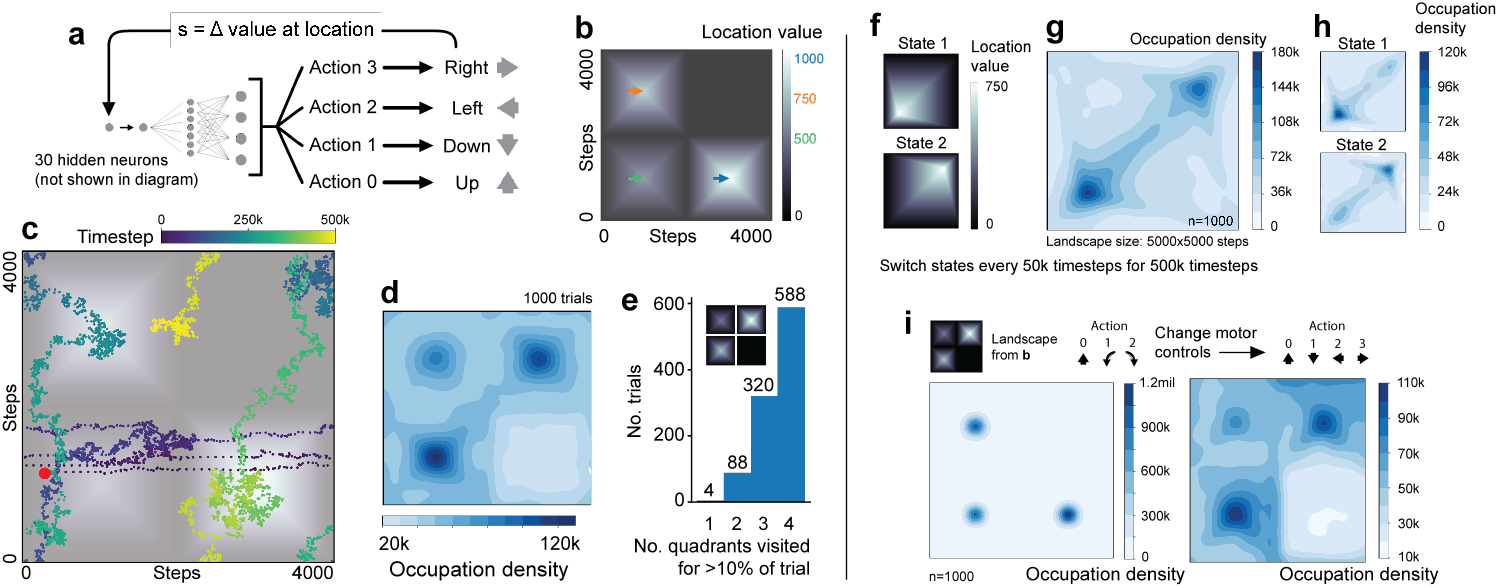
Open-field search task. **a**. Actions correspond to four discrete movements in the reward landscape **b. c**. Every hundredth point for a single trial. Starting point marked in red; see Video 2 for animated track. **d**. Density of occupation for 1000 trials, 500k timesteps each, showing that networks spend more time in higher-value regions. **e**. The number of quadrants occupied for >10% of the trial was counted for each agent. 908 of 1000 agents visited at least 3 quadrants rather than getting stuck at local maxima. **f**. Agents were placed in landscapes that switched between two states every 50k timesteps for 500k timesteps total. **g**. Occupation density over all trials, showing greater occupancy at each maximum. **h**. When (g) is separated by the current landscape state, we see localization at the current maximum. **i**. We change the motor interface of the agent from rotational controls (forward, turn CCW, turn CW) (timesteps 0-500k) to translational controls as in a (timesteps 500k-1mil). Occupation density plots show that agents are able to adapt and continue to seek reward. All experiments in this figure required an upper bound of 2000 CPU hours and 3.5 GB of storage.

##### Agents adapt to changing landscapes

Figure 6f shows a landscape where the reward peak flips between two states every 50k timesteps. The agents distribute their time between the two locations, 6g, and Figure 6h shows that agents localize to the current maximum.

##### PaN can adapt to changes in motor interfaces

Figure 6i places agents in the reward landscape from Figure 6b. For the first 500k timesteps, agents move using a rotation scheme: networks can take three possible actions corresponding to forward movement, 90 degree counterclockwise rotation, or 90 degree clockwise rotation. Then a motor neuron is added and the interface is changed to the standard translational scheme in Figure 6a. Agents seek out higher-reward regions of the landscape throughout the simulation.

### 4.4 PaN autonomously forms task preferences

#### 4.4.1 Task preferences vary between and within networks

PaN displays an autonomy in goal selection that emerges from local interactions. We show this with a network of two input neurons corresponding directly to two output neurons, with one additional output neuron that provides no signal back to the network as a control (Figure 7a). Goal preferences vary over time, shown by plotting preference trajectories, defined as the difference between cumulative Action 1 and Action 0 selections at each timestep. Figure 7b has an example preference trajectory, top, for the action choices below. Figure 7c shows that random seeds can stick to one action for long periods of time, as in the red trace, or they can change preferences often, as in the blue trace. Preferences are distributed between the rewarding actions evenly (Figure 7d).

**Figure 7.**
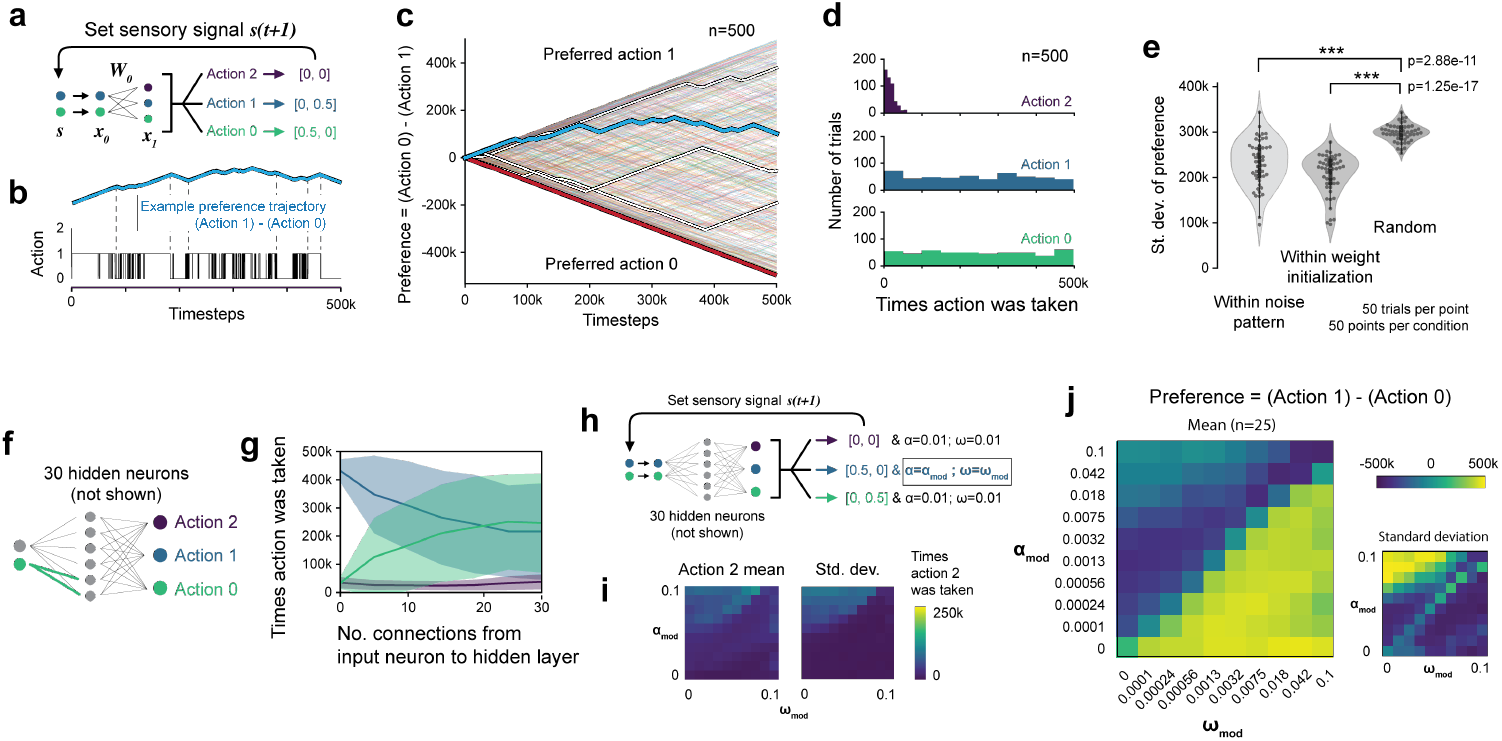
**a**. Network in two-task environments. **b**. Actions can be plotted as a preference trace, defined as the cumulative number of Action 1 selections minus cumulative Action 0 selections. The blue preference trace in b, top, is plotted in **c**. as part of 500 random seeds. Seeds in c. are randomly colored and each trial is 500k timesteps. Several random tracks are highlighted: some networks strongly prefer one action for extended periods of time (red trace) while others change preference rapidly (blue trace). **d**. Networks evenly distribute between Actions 0 and 1 but avoid the non-rewarding Action 2. **e**. Specific noise patterns and weight initializations significantly bias network preference, as measured by the difference between the number of times a network selected Action 1 and Action 0. P-values for two-sided Wilcoxon rank-sum test. **f-g**. We gradually increase the number of connections from the input neuron corresponding to Action 0 and show that the more dense the connections from an input neuron to the hidden layer, the more likely networks will be biased toward the corresponding action. **h**. We change the learning rate for the entire network if Action 1 is selected. Learning rates are by default 0.01 otherwise. **i**. Networks avoid the non-rewarding Action 2. **j**. Networks strongly prefer Action 1 if the modulated activity learning rate α_*mod*_ is lower than the modulated weight learning rate ω_*mod*_. Experiments in figure together required upper bound of 2100 CPU hours and 150 MB for storage.

#### 4.4.2 Task preferences can be biased in biologically relevant ways

##### Noise and initialization

We plot the standard deviations of preference within noise patterns for differently initialized networks in Figure 7e, left, as well as standard deviations of preference within the same weight initializations using different noise patterns in Figure 7e, center. We compare both cases to random noise patterns and initializations in Figure 7e, right, showing that preferences are biased by specific patterns of activity and weight noise.

##### Connectivity

In networks with two input neurons and 30 hidden neurons, we increase the number of connections from the input neuron corresponding to Action 0 and keep the other input neuron fully connected (Figure 7f). The sparser the connectivity of the input neuron, the less the network prefers its corresponding action (Figure 7g).

##### Modulated learning rates

Next, we change the activity learning rates α and weight learning rates ω when networks choose Action 1 (see Algorithm 1 for variable definitions and Figure 7h for illustration). Networks retain their reward-seeking behavior by avoiding the non-rewarding Action 2 (Figure 7i). But interestingly, Figure 7j shows that when the modulated activity rate α_*mod*_ is less than the modulated weight learning rate ω_*mod*_, networks strongly favor the action responsible for the change (Action 1). In other words, if an action results in faster weight learning than activity learning, then that action is preferred (see [9] for biological plausibility).

#### 4.4.3 Open-field search task with two types of reward

We then evaluate whether we can bias PaN’s preferences in the open-field search task. We add a second input neuron to the network in Figure 6a, shown in Figure 8a, and allow it to freely behave in the landscape in Figure 8b. Without any bias, networks do not consistently prefer either maximum over the other (Figure 8c). When we bias networks toward Landscape II by decreasing connections from the input neuron corresponding to Landscape I (Figure 8d), networks prefer the Landscape II maximum (Figure 8e-f). When we use the learning rate modulation method to bias the network, decreasing the activity learning rate for signals from the biased landscape (Figure 8g), we find that networks can form extremely strong preferences for the currently biased landscape. Preferences were rapidly switched by changing the biased landscape every 50k timesteps for 500k timesteps total (Figure 8h), with an example track in Figure 8i.

**Figure 8.**
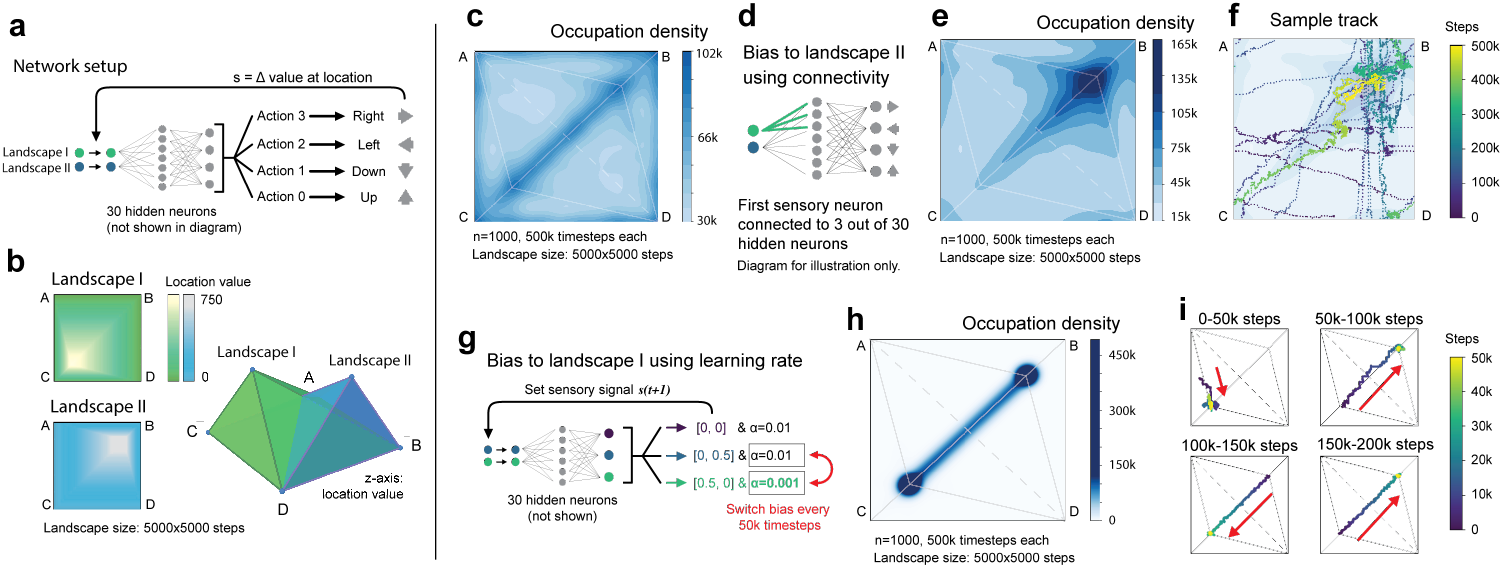
**a**. Networks can take translational movements in **b**. a landscape with two overlapping value gradients. **c**. An unbiased network chooses either of the value maxima. **d**. When networks are biased by reducing the connections from the input neuron corresponding to Landscape I values, **e**. networks overwhelmingly prefer Landscape II. **f**. Sample track for e. **g**. Next we bias networks by reducing activity learning rates when they receive positive Landscape I signal. Every 50k timesteps, the bias switches to the other landscape for a total of 500k timesteps. **h**. Networks move between landscape maxima, and a sample track in **i**. shows that networks move quickly toward the biased landscape, with arrows to emphasize direction of movement. Experiments in this figure required upper bound of 1500 CPU hours and 3 GB.

## 5 Discussion

We show that neural networks using only local prediction and noise can exhibit flexible, emergent, and autonomous behavior. Our algorithm PaN can switch between exploration and exploitation without external prompting, and these rich behaviors are explained by attractor dynamics. PaN is distinct from classical RL agents in that it uses entirely local predictive updates rather than an environmental reward function; consequently, it has the potential to select and shape its own goals. We show that PaN networks can have preferences and can change them over time, similar to observations in animals (see fruit fly behavior in [40]). And because of how PaN differs from other algorithms that learn through environmental interaction, we define a preliminary framework called *self-supervised behavior* (SSB) and elaborate upon it in Appendix A.

Our algorithm is biologically plausible, has the potential to account for autonomy in living nervous systems, and shows that emergent behavior is capable of switching between exploration and exploitation, where reward signals can be maximized during exploitation. In our estimation, these features make PaN a possible starting point for more expressive algorithms that better resemble natural intelligence.

### 5.1 Limitations

We have only tested PaN in simple environments. We have not demonstrated PaN’s use of some of the advantages of predictive coding models found in the literature [41], such as their ability to learn useful sensory representations; nor have we shown any capacity of PaN to form associative memories, which are required in more complicated tasks, or how aversive stimuli might be represented. Moreover, deeper PaN networks, ones with recurrent connections, or networks following Dale’s Law [10] may allow for more complex behavioral dynamics that PaN is incapable of in its current form. These are all important missing components and will be interesting directions for future work.

## Acknowledgments and Disclosure of Funding

We thank members of the Kreiman and Ramanathan Labs as well as Benjamin de Bivort, Michael Brenner, and Kristopher T. Jensen for discussions and comments. Support was provided by NIH 5R01NS117908-03 (SR), NIH-R01EY026025 (GK), the Fetzer Foundation (GK), and the NSF GRFP Fellowship (CL). No competing interests to declare.

## A Self-supervised behavior: a novel framework for environmental interaction

At the moment, reinforcement learning (RL) is the primary framework for artificial agents that learn from and act within an environment. However, critics have pointed out that RL may be limited, in part because it requires the foreknowledge of an appropriate goal [24]. While we do define a single predictive loss for PaN (Equation 1), the loss is determined internally and updates include noise. This is in contrast with RL, where learning is set up as a way to maximize expected return from external Markov Decision Processes [43]. We argue that our algorithm is distinct both in its definitions and resultant behaviors, which are never designed to achieve an optimal action trajectory (the authors of GFlowNets make a similar argument to distinguish their work from RL; see [3]).

PaN is only a preliminary model, but we believe it has the potential to be extended to autopoietic computational systems. Most modern machine learning algorithms cannot by definition [24], because they use backpropagation to learn and backpropagation requires tasks or objectives that are predefined. In self-organized, self-managing, and scalable systems [5], however, goals should emerge on their own. We therefore suggest that PaN belongs to a new framework, which we call self-supervised behavior (SSB), named because PaN’s energy function resembles that of models that use self-supervision over time [35] to learn sensory representations. Here we posit features that an SSB model should possess.

### A.1 Features of self-supervised behavior

- **Updates are based on internal, local loss, with the absence of an environmentally defined global utility function**. An SSB agent should exhibit emergent goals and behaviors rather than ones that are predefined. Emergence allows for diversity in behavior both between and within individual agents, as well as a great capacity to adapt to changing circumstances.
- **Autonomous operation that can execute continual learning in dynamic situations**. SSB should remain autonomous. For an agent to function—much like a living organism—it should not require continual direction from an external entity. An SSB agent’s ability to carry out continual learning should be entirely self-driven.
- **Evidence of apparent goals that are self-selected, and evidence of ability to improve performance on these goals**. Again, goal selection should emerge from an SSB agent’s internal workings rather than an external definition. The agent should also be able to learn over time to better achieve these goals, whether through increasing a reward signal, which we have shown here, or by learning effective motor sequences on its own, which we leave for future work.

### A.2 Why define an alternative framework?

Defining SSB allows us to scaffold ideas and results from PaN without relying too much on the details of its implementation. We do not believe PaN’s particular energy function, architecture, and training methods are unique in their ability to produce the results we show in this paper. At the same time, we believe PaN’s use of emergence to generate behaviors makes it fundamentally unlike most other algorithms.

Namely, we feel it is important to define a framework for agents behaving in environments that *does not* rely on achieving some optimal course of action. RL and even active inference begin from the assumption that actions taken in an environment serve to optimize some set task or environmental function (minimizing overall “surprise” in active inference [36], for example). But one might consider the great diversity, creativity, and self-driven nature of animal behavior, and that all of it might suggest that learning algorithms do not need to optimize an environmental outcome (outside of fulfilling the basic constraints of survival and reproduction) in order to perform useful, adaptive, or simply interesting behaviors.

Thus we believe that defining SSB as a framework is important and ongoing work, for PaN and other emergent algorithms. Our hope is that suggesting this alternate framework for agents acting in environments will encourage researchers to consider that explicit optimization is only one way of learning from an environment, and to explore other, possibly richer ways of interacting with the world.

## B Explore/exploit modes persist in entropy calculations with different window sizes

In Figure 9 we show that distinct explore/exploit states persist regardless of window sizes for entropy calculations. The data here are for the system in Figure 10. First we plot comparisons for PaN (Figure 9a) and an ϵ-greedy agent with matched reward (Figure 9b). Figures 9c-d show PaN at different timescales, while Figure 9e plots the entropy for a 1000-timestep rolling window for both PaN and ϵ-greedy agents. PaN exhibits distinct high-entropy (explore) and low-entropy (exploit) phases, while ϵ-greedy agents maintain consistent levels of exploration throughout. Figure 9f shows that even when entropy windows are very small, PaN still has a bimodal entropy distribution, while the ϵ-greedy agent distributions are unimodal. Figure 9g plots exploitation, as defined by the frequency of windows with very low entropy (marked bars in Figure 9f) for a range of noise conditions.

**Figure 9.**
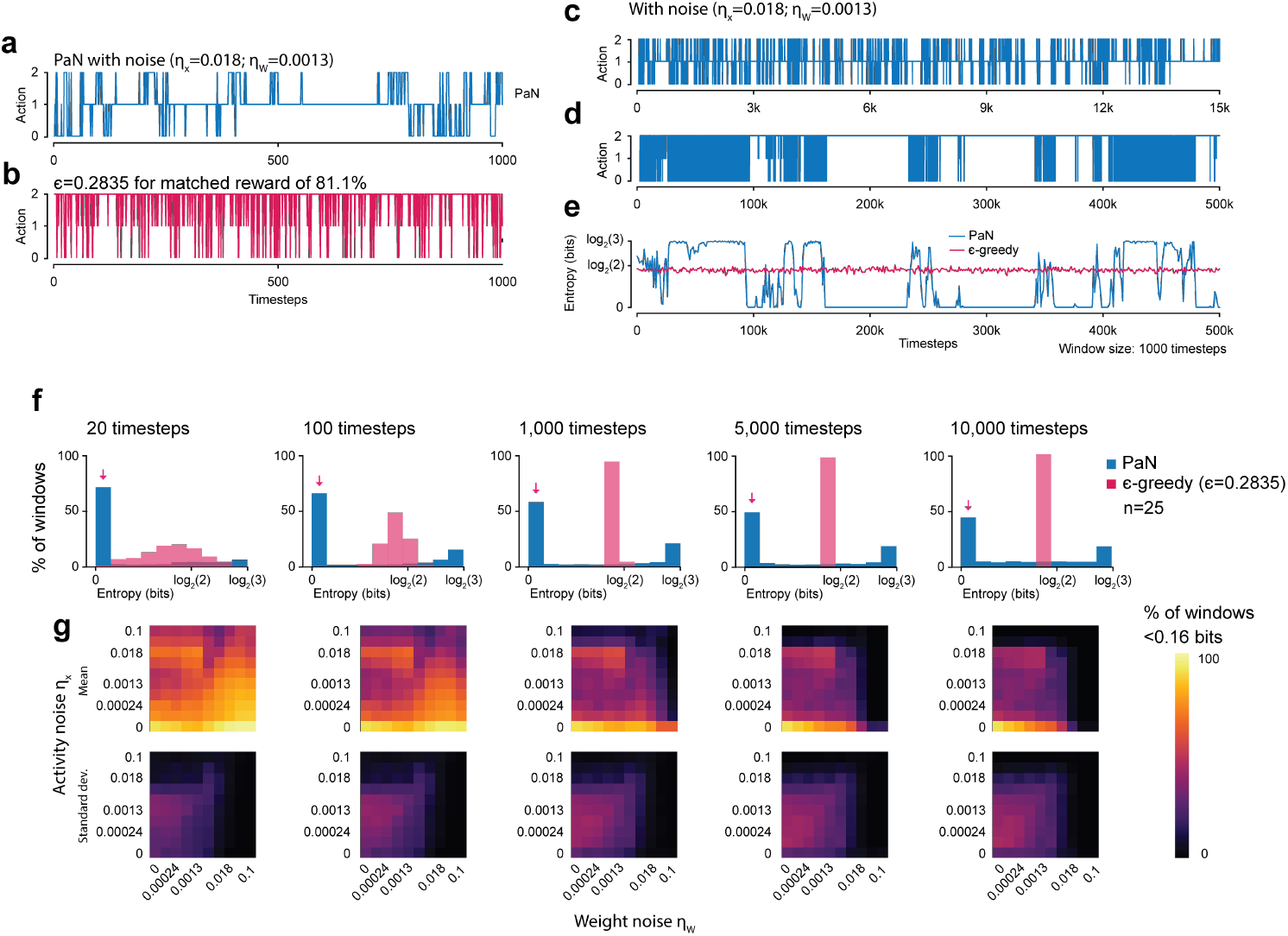
**a**. Sample action choices for a PaN network in Figure 10a, first 1000 steps. Compare to **b**., which is an ϵ-greedy agent with a matched proportion of reward selected. **c**. The first 15k steps from the same trial as a. PaN displays exploration (dense parts of plot) and exploration (fixation) at multiple timescales. **d**. Entropy of rolling windows of action selections for both PaN and ϵ-greedy agents. PaN transitions between high entropy (exploration) and low entropy (exploitation). **e**. Histograms of entropies for PaN and ϵ-greedy agents with differently sized entropy windows, showing that bimodality in PaN’s entropy does not depend strongly on window size. Here bimodality suggests the presence of two behavioral states, as discussed for Figure 3h in main text. **f**. We devise a proxy to visualize the amount of exploitation in a PaN network, which is the proportion of windows across trials and 25 random seeds with < 0.16 bits (0.1 * log_2_(3)) of information. Pink arrows mark this value in e. for a noise setting of *η*_*x*_ = 0.018; *η*_*W*_ = 0.0013. We plot the exploitation metric for all noise settings and a range of window sizes. For a broad range of window sizes (from 1k to 10k timesteps tested here) there remains a region of noise that allows for exploitation, around *η*_*x*_ = 0.018.

**Figure 10.**
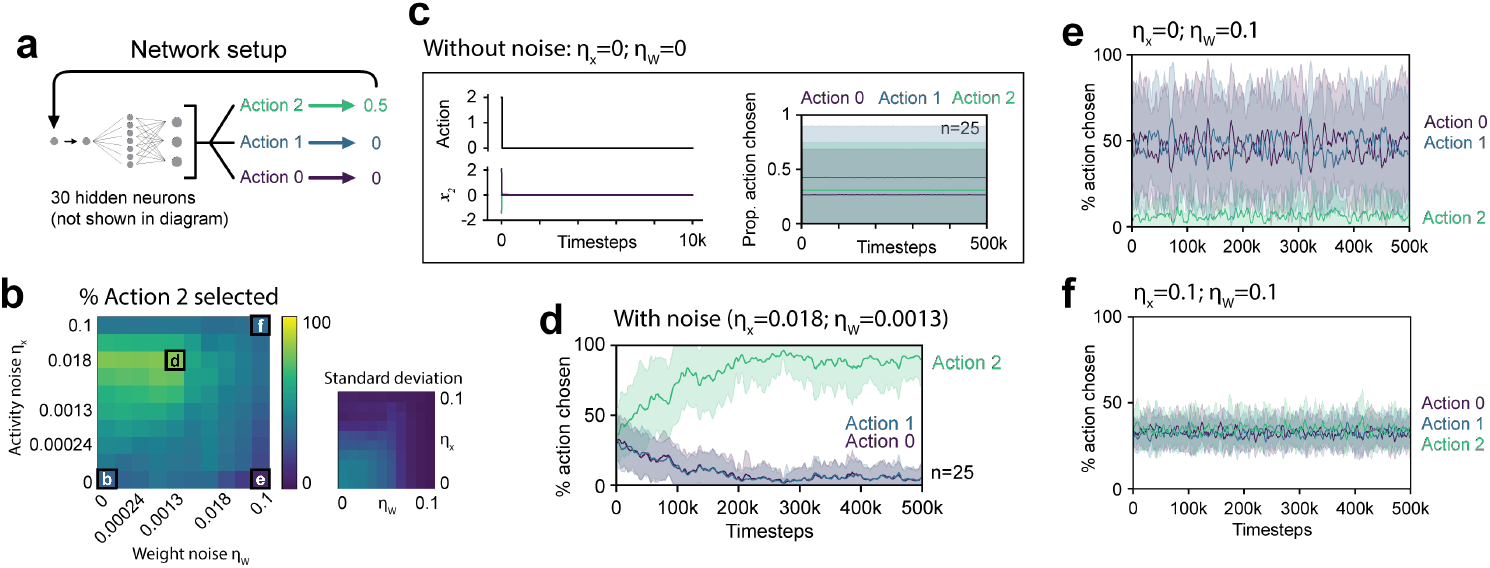
PaN behavior in a 3-armed bandit task under different noise settings. **a**. Network setup with one hidden layer and three possible actions. Actions correspond to [0, 0, 0.5] reward, respectively. **b**. The percentage of actions chosen that were rewarding (Action 2). Mean (left) and standard deviation (right); n=25 per condition. For exact values see Table 2. **c**. When noise parameters are 0, networks fixate on one randomly chosen action. This and following plots are mean action selections over time, with shaded areas denoting standard deviation, n=25. **d**. When noise is nonzero and within a certain range (see b.), networks learn to choose the most rewarding action. **e**. When weight noise is very high and activity noise is 0, rewarding actions (Action 2) are *not* preferred, and the network chooses the zero-reward actions preferentially. **f**. In high noise settings, networks are dominated by noisy actions throughout trials.

## C The effect of noise parameters on behavior

The scales of activity and weight noise affects how frequently a PaN network chooses the rewarding action in a bandit task. They also affect the amount of exploration, and variability in performance between networks. Here we show, using a network with a 30-neuron hidden layer and 3 possible actions, that different noise settings can produce different qualitative regimes of behavior (Figure 10).

**Table 2:** Values for Figure 10b. **a**. Mean and **b**. standard deviation of proportion of actions taken that were rewarding for an action space with 3 actions corresponding to rewards [0.5, 0, 0]. For three-neuron system in Figure 10a. Note that here, random actions would correspond to a mean of 0.333. The maximum mean in (a) is bolded (.811).

| $\eta_x \backslash \eta_W$ | 0 | .0001 | .00024 | .00056 | .0013 | .0032 | .0075 | .018 | .042 | .1 |
| --- | --- | --- | --- | --- | --- | --- | --- | --- | --- | --- |
| 0 | .307 | .319 | .339 | .386 | .359 | .343 | .340 | .245 | .173 | .063 |
| .0001 | .401 | .359 | .372 | .401 | .381 | .373 | .424 | .366 | .317 | .211 |
| .00024 | .464 | .450 | .408 | .427 | .401 | .383 | .430 | .373 | .332 | .236 |
| .00056 | .507 | .534 | .515 | .497 | .454 | .402 | .440 | .382 | .348 | .264 |
| .0013 | .604 | .621 | .598 | .595 | .547 | .442 | .459 | .397 | .365 | .297 |
| .0032 | .636 | .651 | .662 | .693 | .670 | .541 | .500 | .423 | .389 | .335 |
| .0075 | .717 | .732 | .728 | .768 | .757 | .556 | .553 | .460 | .411 | .361 |
| .018 | .800 | .796 | .789 | .802 | **.811** | .587 | .516 | .482 | .416 | .359 |
| .042 | .645 | .638 | .634 | .635 | .622 | .541 | .420 | .439 | .392 | .347 |
| .1 | .387 | .386 | .382 | .390 | .388 | .388 | .352 | .352 | .364 | .346 |

(b)
| $\eta_x \backslash \eta_W$ | 0 | .0001 | .00024 | .00056 | .0013 | .0032 | .0075 | .018 | .042 | .1 |
| --- | --- | --- | --- | --- | --- | --- | --- | --- | --- | --- |
| 0 | .441 | .412 | .401 | .361 | .305 | .264 | .155 | .066 | .030 | .012 |
| .0001 | .418 | .402 | .397 | .363 | .306 | .260 | .147 | .054 | .026 | .013 |
| .00024 | .408 | .411 | .391 | .356 | .305 | .256 | .145 | .055 | .026 | .013 |
| .00056 | .351 | .366 | .348 | .330 | .293 | .242 | .142 | .054 | .026 | .012 |
| .0013 | .273 | .271 | .269 | .284 | .255 | .208 | .136 | .053 | .025 | .013 |
| .0032 | .242 | .232 | .220 | .200 | .212 | .188 | .125 | .053 | .025 | .011 |
| .0075 | .128 | .123 | .133 | .133 | .144 | .168 | .119 | .054 | .024 | .010 |
| .018 | .049 | .065 | .063 | .071 | .072 | .145 | .114 | .050 | .020 | .010 |
| .042 | .067 | .070 | .088 | .081 | .090 | .134 | .075 | .040 | .019 | .008 |
| .1 | .047 | .051 | .045 | .044 | .048 | .040 | .036 | .035 | .024 | .010 |

In Figure 10a we show the network setup and in Figure 10b, we plot the percentage of time the rewarding action was selected. We see that there is a range of noise settings that leads to higher reward, around *η*_*x*_ *∈* [0.0013, 0.042]. Without noise, networks fixate on a random action in Figure 10c. With noise, networks choose the rewarding action and learn to exploit it more over time (Figure 10d).

An interesting case arises, however, when weight noise is high and activity noise is low. We see in Figure 10e that here, the rewarding action is avoided rather than preferred. This is because noise is only propagated through the system when there is signal, and so the network prefers zero reward actions over ones with any signal.

Finally, when weight and activity noise are both high, all actions are random (Figure 10f).

### C.1 Table values for noise parameter sweeps

We include here the values for heatmaps in Figure 3i-j and Figure 10b.

## D Analysis of attractor dynamics

The goal of this section is to develop an intuition and explanation for PaN behavior based on a dynamical systems perspective of the analytically tractable two-neuron network. We begin by reinterpreting the prediction and noise (PaN) algorithm as a system of stochastic differential equations (SDEs). We use these equations to derive an attractor in the network’s parameter space in the absence of noise. The attractor has three components and we describe their associated stabilities and behaviors. Finally, we simulate the long-term distributions of network parameters along the attractor, and show how these distributions change with different noise magnitudes.

### D.1 The general PaN algorithm as a set of difference equations

PaN’s dynamics are governed by the predictive energy function (Section 3):

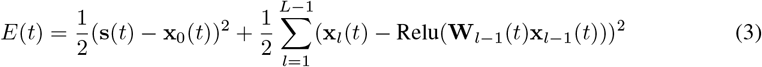

At timestep *t*, a network’s activities **x**_*l*_(*t*) and weights **W**_*l*_(*t*) are updated according to the following steps, which we rewrite from Algorithm 1. Here, α and ω are the activity and weight learning rates, s_*x*_ and s_*W*_ are the activity and weight noise standard deviations, and *η*_**x***/***W**_(*t*) are i.i.d. standard normal noise across parameters and time (notation here differs slightly from Algorithm 1 to facilitate analysis).

1. Begin with activities and weights (**x**_*l*_(*t*), **W**_*l*_(*t*)) at time *t*.
2. Perform gradient descent on *E* with respect to the activities **x**_*l*_

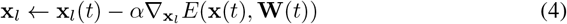
3. Add activity noise

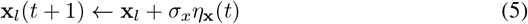
4. Perform gradient descent on *E* with respect to the weights **W**_*l*_

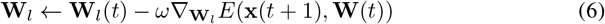
5. Add weight noise

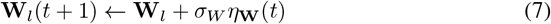

We can summarize the change in the activities and weights between timesteps *t* and *t* + 1, Δ**x**_*l*_(*t*) and Δ**W**_*l*_(*t*), as

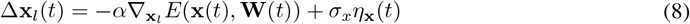

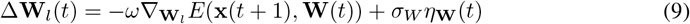

Notice that Δ**W**_*l*_(*t*) depends on **x**(*t* + 1) rather than **x**(*t*). So the weight update at timestep *t*, written in terms of the parameters at that timestep, is given by

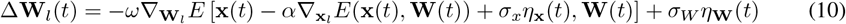

We can summarize the PaN algorithm with the following set of noisy difference equations

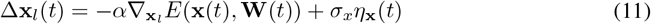

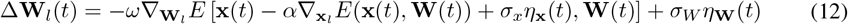

From now on, we drop the *t* argument, since it is the same everywhere.

### D.2 The two-neuron PaN network as a noisy dynamical system

#### D.2.1 Difference equations for the two-neuron network

We now focus on the dynamics of the two-neuron agent in the two-action bandit task. The agent is parameterized by two activities, *x*_0_ and *x*_1_, and a weight *W*. It receives one sensory input *s* from the environment. Its dynamics are governed by the energy function from Equation 2:

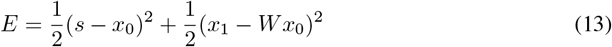

Here we removed the ReLu nonlinearity from Equation 1 for ease of analysis. Following the previous section, this energy gives the update rules

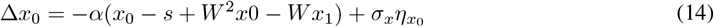

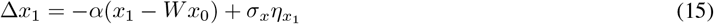

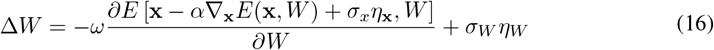

The activity noise now appears in the weight updates, meaning that activity noise and weight updates are correlated. Upon expansion of Δ*W*, one finds that the variance of the noise in the weight update also depends on *x*_0_, *x*_1_, and *W*. Both of these observations will complicate future analysis.

Luckily, we can make a simplification: the complexity of the expression for Δ*W* comes from the additional − α *∇*_**x**_*E*(**x**, *W*) + σ_*x*_*η*_**x**_ term. This term appeared because we updated the weights according to **x**(*t*+1) rather than **x**(*t*). However, terms from this addition must be of *O*(ωα), *O*(ωσ_*x*_), or higher in α or σ_*x*_. In all our simulations, these three parameters are small (on the order of 10^−2^ or less), so we neglect the higher-order terms. In Section D.4, we empirically show the validity of the approximation, which gives Δ*W* the much simpler approximate form

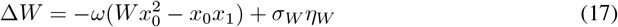

Because PaN interacts dynamically with its environment, the sensory stimulus it receives is a function of its motor output *x*_1_. Indeed, as per the definition of the two-lever bandit task, *s* is given by

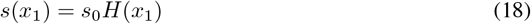

where *H*(*x*) is the Heaviside step function and *s*_0_ is the value of the reward lever. ^1^ We can rewrite two-neuron PaN dynamics as the following system of difference equations:

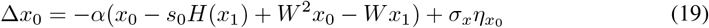

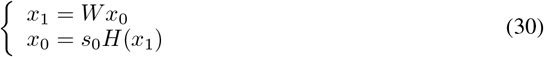

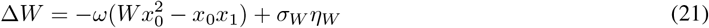

This is the form of a discretized system of stochastic differential equations (SDEs).

#### D.2.2 Stochastic differential equation approximation for the two-neuron system

The three noise terms are now uncorrelated normal random variables, making the equations above amenable to analysis. We now demonstrate that the PaN algorithm can be interpreted as an SDE, giving insight into the long-term parameter distributions we should expect.

A general SDE takes the form:

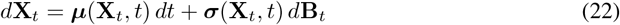

Here, **X**_*t*_ and ***µ***(**X**_*t*_, *t*) are vectors in ℝ^*N*^. ***s***(**X**_*t*_, *t*) is an *N × M* matrix, and **B**_*t*_ is an *M*-dimensional Wiener process. If we discretize time in steps Δ*t*, the Euler scheme implies

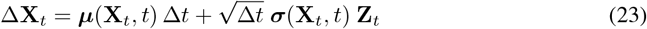

where Δ**X**_*t*_ = **X**_*t*+Δ*t*_ − **X**_*t*_ and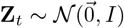. Letting

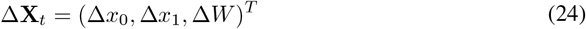

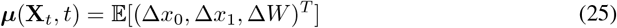

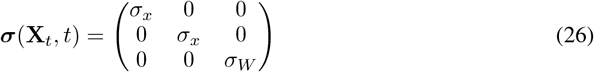

we see that the PaN is a 3-dimensional SDE discretized by the Euler scheme with Δ*t* = 1. In Section D.4 we support this claim by showing that the system’s long term behavior is consistent with the real PaN agent as we send Δ*t* → 0 in simulations.

This result motivates us to reinterpret the two-neuron PaN agent, and all PaN systems in general, as SDEs with nonlinear drift terms and diagonal σ matrices. Note that the form of the SDE depends both on the architecture of the agent through *E*(*t*) and on the environment function through the form of *s*(**x**). This interpretation holds even in stochastic environments where the stimulus vector **s** is not completely determined by the network activities **x**, as we can always approximate it as *s*(**x**) plus an additional noise term.

For the linear two-neuron PaN agent in the deterministic bandit task, the SDE that describes the system is given by

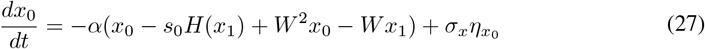

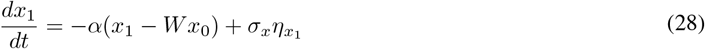

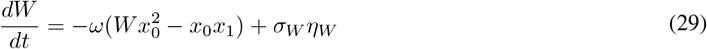

where here the *η* denote independent white noise processes.

### D.3 Derivation of attractors and how they correspond to explore or exploit behavior

We can gain a geometric intuition for PaN behavior by considering the attractor of the dynamical system (the set of all points that are stable without noise). To find the attractor, we calculate the fixed points and determine stabilities with linear stability analysis.

#### D.3.1 Linear stability analysis of the fixed points

Ignoring noise and setting all derivatives to zero, we find that the fixed points are given by

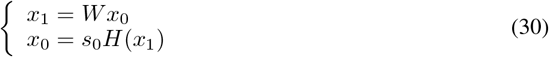

We perform linear stability analysis to determine which fixed points form an attractor. Letting 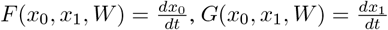, and 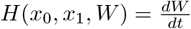, the Jacobian is given by

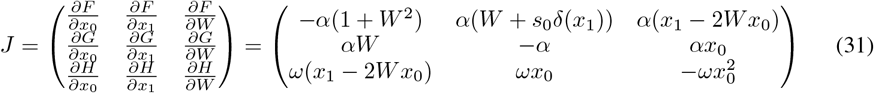

Since 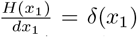, where d(*x*_1_) is the Dirac delta function. The Dirac delta distinguishes two cases, *x*_1_ ≠ 0 and *x*_1_ = 0.

**Stability for fixed points where** *x*_1_ ≠ 0 We begin by analyzing the system’s stability when *x*_1_ ≠ 0. This occurs if and only if *x*_0_ ≠ 0 and *W* ≠ 0. At the fixed points, if *x*_0_ ≠ 0, then *x*_0_ = *s*_0_. So, we evaluate the Jacobian where *x*_0_ = *s*_0_ and 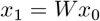 and solve for its eigenvalues λ. We find

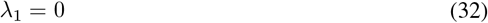

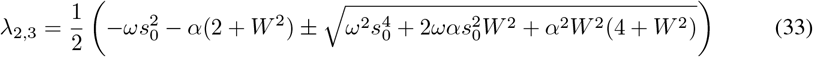

The zero eigenvalue corresponds to the line of fixed points. Since α > 0 and ω > 0, the second two eigenvalues are negative real numbers for all *W*. As such, the fixed points where *x*_1_ ≠ 0 are all stable.

**Stability for fixed points where** *x*_1_ = 0 Next, we analyze the case where *x*_1_ = 0. Strictly speaking, the Jacobian is undefined here because δ(*x*_1_ = 0) =. However, we momentarily consider the error function approximation of *H*(*x*_1_) so that its derivative is not δ(*x*_1_) but rather a very sharp Gaussian. This means we can approximate our Jacobian by replacing δ(0) with Ω, where Ω is an arbitrarily large but finite positive number.

When *x*_1_ = 0, there are two sub-cases. Either *x*_0_ = 0 (a line where *W* can take any value) or *x*_0_ = *s*_0_ and *W* = 0 (a point). With the Ω approximation, the eigenvalues of the Jacobian on the line where *x*_0_ = 0 are

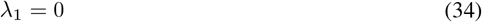

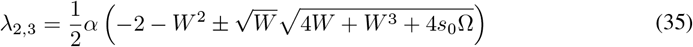

The eigenvalues at the fixed point characterized by *x*_0_ = *s*_0_ and *W* = 0 are

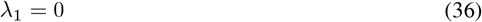

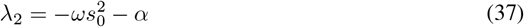

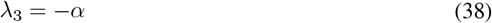

In Equations 36-38, all nonzero eigenvalues are negative, so this fixed point is stable without noise. However, Equation 35 has an eigenvalue with a positive real part when 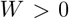. This is because Ω >> 1, so if the term in the square root is multiplied by a positive real value 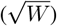, then one of the eigenvalues will be a very large positive number.

We therefore conclude that the fixed points where *W* > 0 and *x*_0_ = 0 are not stable. However, all other fixed points where *x*_1_ = 0 are stable without noise.

#### D.3.2 The attractor of the two-neuron network

Combining the results of Section D.3.1, we find that there is an attractor in the two-neuron network’s parameter space given by

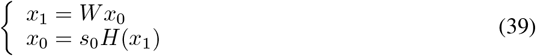

with the constraint that *W ≤* 0 if *x*_0_ = 0. We visualize it in Figure 11—it looks like two disconnected line attractors in the parameter space of (*x*_0_, *x*_1_, *W*).

**Figure 11.**
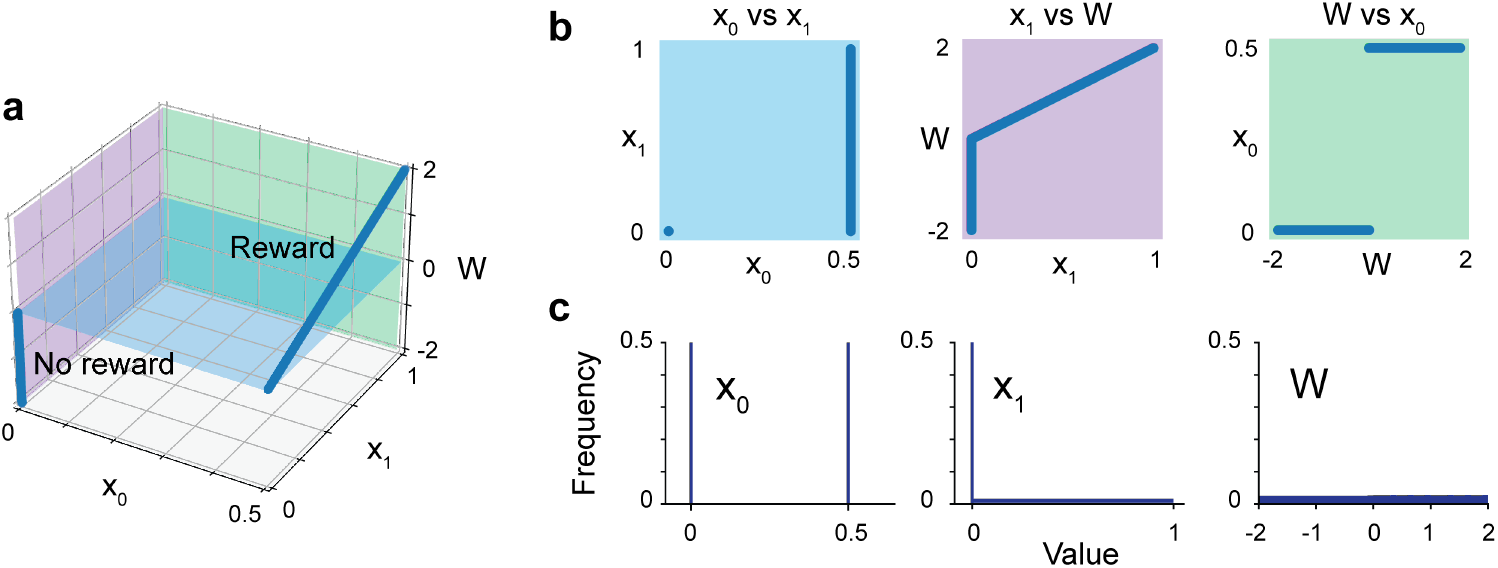
Attractors without noise in the two-neuron network. **a**. Reward and no-reward attractors plotted in the parameter space of (*x*_0_, *x*_1_, *W*). **b**. Projections of attractors on each labelled plane, color-matched to planes in a. **c**. Frequency distributions of each parameter.

#### D.3.3 Without noise, all parts of the attractor are stable

Projected into the three planes, the attractor looks as in Figure 11b, with distributions of each parameter in Figure 11c. The histograms give the expected distribution of the agent’s parameters under the zero noise condition: networks stay on the attractor and do not favor any section of the attractor over another. These plots can be used as references for later figures.

#### D.3.4 With motor noise, only the rewarding part of the attractor is stable

If we add noise, any part of the attractor that lies on the agent’s “decision boundary,” *x*_1_ = 0, is unstable. This is because when *x*_1_ = 0, small amounts of noise added to *x*_1_ can switch the agent’s lever choice, which alters whether PaN is driven to the reward or no reward attractor. The entire no reward attractor lies on this decision boundary, which means that it is unstable in the presence of motor noise.

The reward attractor, on the other hand, is almost entirely stable. Only the very edge (where *x*_1_ = *W* = 0) lies on the decision boundary, so only when PaN diffuses along the attractor to the edge can it select the zero reward lever and begin converging to the zero reward attractor.

We show these results in Figure 12. When there is no motor noise as in Figure 12a-f, networks do not prefer the rewarding action, best seen in the *x*_0_ distributions in Figures 12d, f, which are evenly split between inputs 0 and 0.5. When motor noise is added as in Figure 12g-j, networks do prefer the rewarding action. This is again best seen in the *x*_0_ distributions in Figures 12h, j, where now the 0.5 input is strongly preferred.

**Figure 12.**
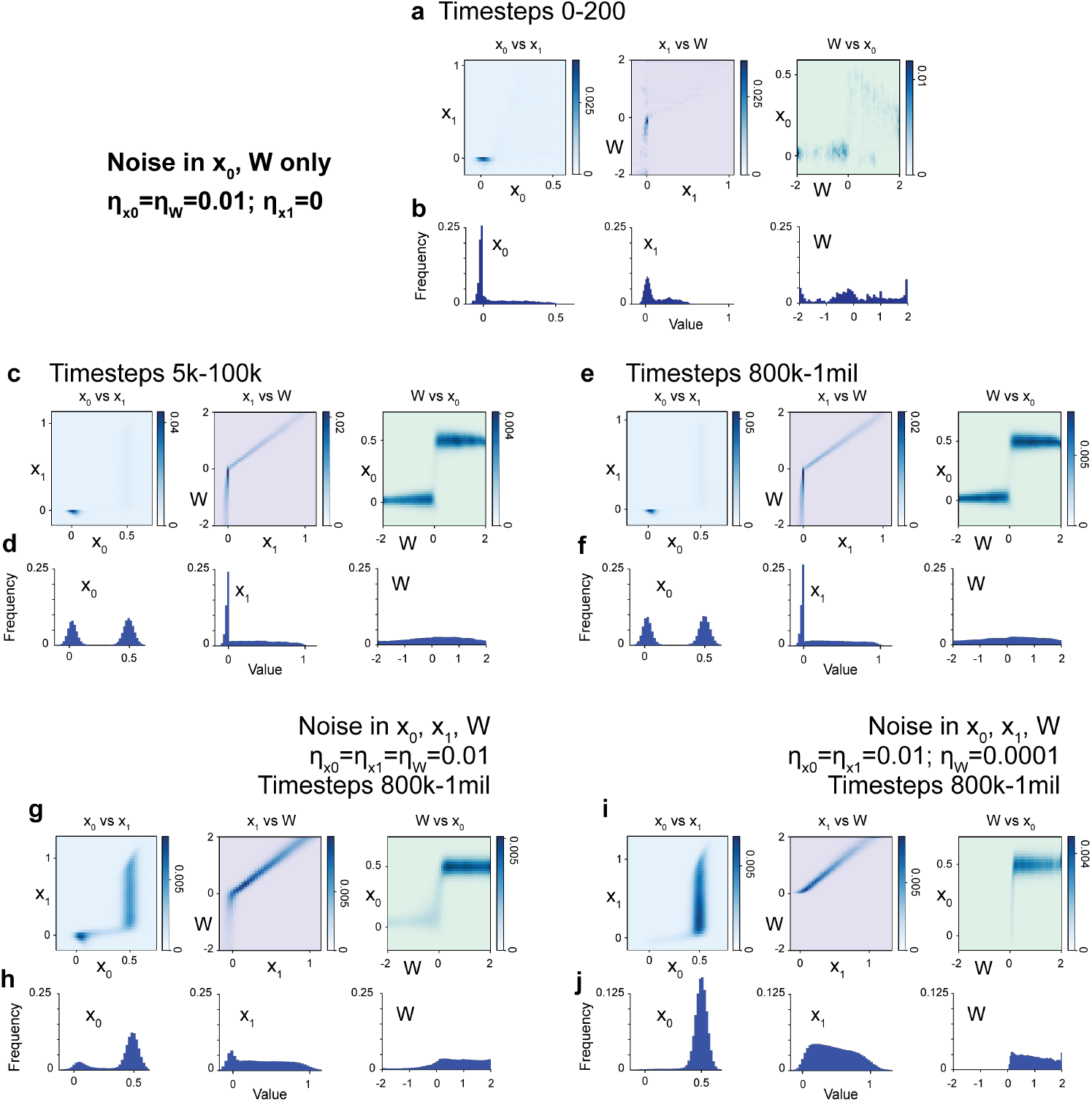
Motor noise (noise at *x*_1_) is necessary for reward-seeking behavior. **a-f** look at networks without motor noise,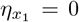. Networks are still settling in the first 200 timesteps (a-b) but by timesteps 5k-100k (c-d), they have settled into their long-term distributions (e-f). Networks evenly distribute between choosing both actions, most easily seen in the *x*_0_ histograms in (d) and (f). **g**. With motor noise, networks prefer the rewarding action, which is again best seen in the *x*_0_ distribution in **h. i-j**. Noise can be tuned to further push networks toward exploitation. With less weight noise, networks almost entirely choose the rewarding action (j).

#### D.3.5 Attractor dynamics explain overall explore/exploit behavior

Given the observations up until now, we expect the following behavior.

1. The agent should in general be converging to either the reward or no reward attractor, depending on its current lever choice.
2. In the presence of noise, the agent should spend more time at the reward attractor because it is more stable.
3. Since the agent can switch from converging towards the reward attractor to converging towards the zero reward attractor when it is near the reward attractor edge, *x*_1_ = *W* = 0, there should be an occasionally traversed “bridge” along the *x*_0_ axis between the no reward and reward attractors.

Now we have a geometric explanation for the agent’s long-term explore-exploit behavior. We can interpret the time that the agent spends diffusing along the reward attractor as its exploit phase. This phase should be longer than all other phases because the reward attractor is the most stable section of the agent’s parameter space. We can interpret the time that the agent spends switching between the two attractors, along the *x*_1_ = *W* = 0 “bridge”, as its explore phase. This phase should happen whenever the agent nears the edge of the reward attractor.

The rate at which the agent oscillates between explore and exploit phases therefore depends on the length of the reward attractor, its rate of diffusion along the attractor (which is proportional to the noise in the system), and the distance between the reward and zero reward attractors, which depends on the magnitude of *s*_0_. Quantifying these rates will be interesting directions for future work.

With this picture, we can also explain why it takes time for the agent to learn to maximize reward. The learning phase is the transient phase that occurs while the agent converges to an attractor from its random initial point, which one can see in Video 1.

### D.4 Empirical validation of attractor interpretation under various noise conditions

Here we show that the approximations made in Section D.2.1, where higher-order terms are dropped in the difference equations, do not have a noticeable effect on network distributions. All plots in Figure 13 have noise in all three parameters: 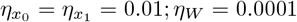. Plots are all for timesteps 300k-500k. Figure 13a-b plot parameter projections and distributions when higher-order terms are included. There is little difference between Figure 13a-b and Figure 13c-d, when higher-order terms are dropped, suggesting that dropping these terms is a valid approximation.

**Figure 13.**
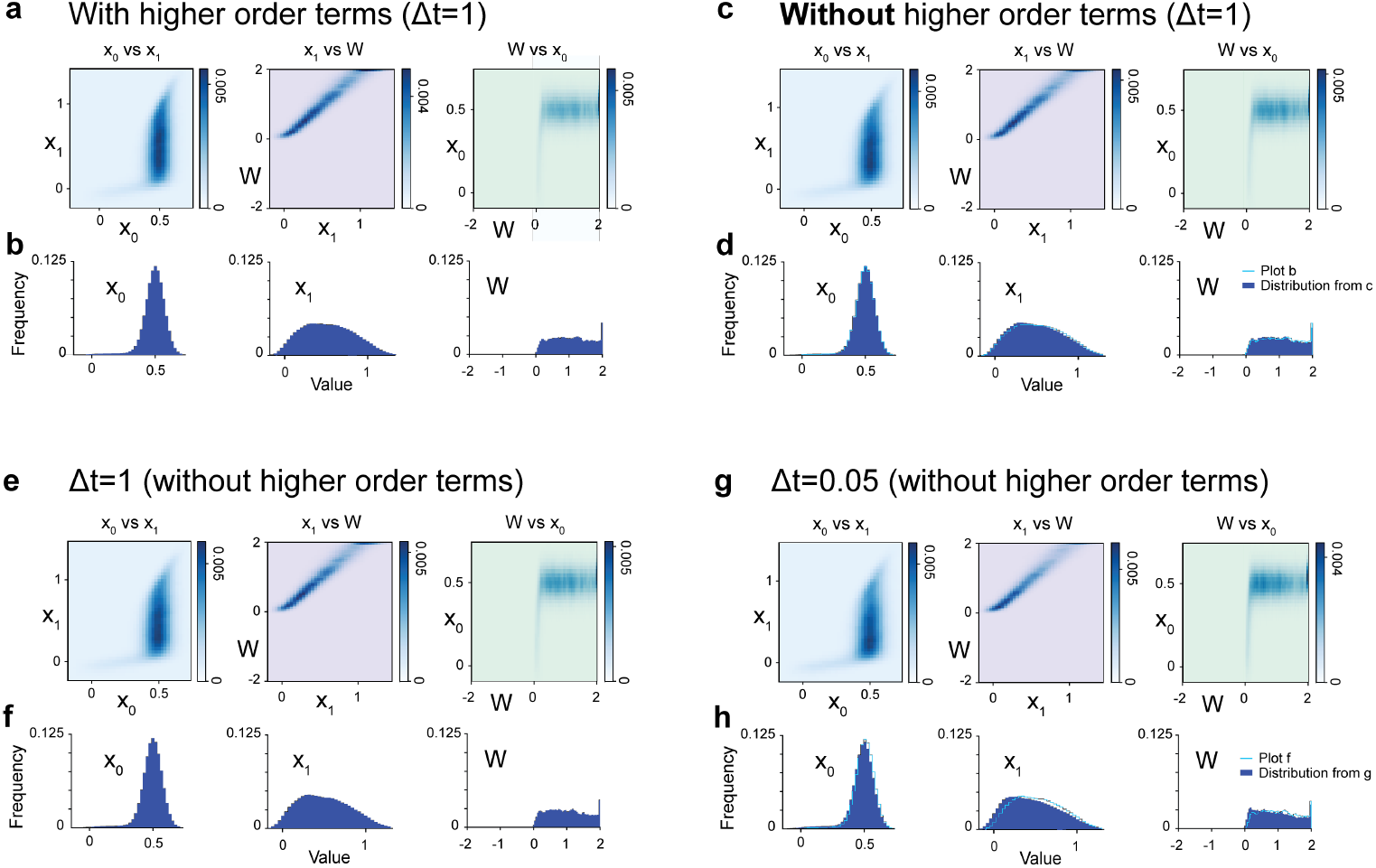
Support for approximations made in Sections D.2.1 and D.2.2. All plots are for two-neuron PaN networks with noise in all three parameters: 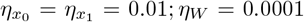. Plots are all for timesteps 300k-500k. **a-b**. Projections and distributions, respectively, for parameters when higher-order terms are included in the difference equations in D.2.1. **c-d**. Projections and distributions when higher-order terms are dropped. Differences between a-b. and c-d. are very slight. **e-f**. Timestep of 1 as in Section D.2.2, compared to a timestep of 0.05 as in **g-h**. There is again very little difference, suggesting that PaN can be written and understood as a discretized SDE.

**Figure 14.**
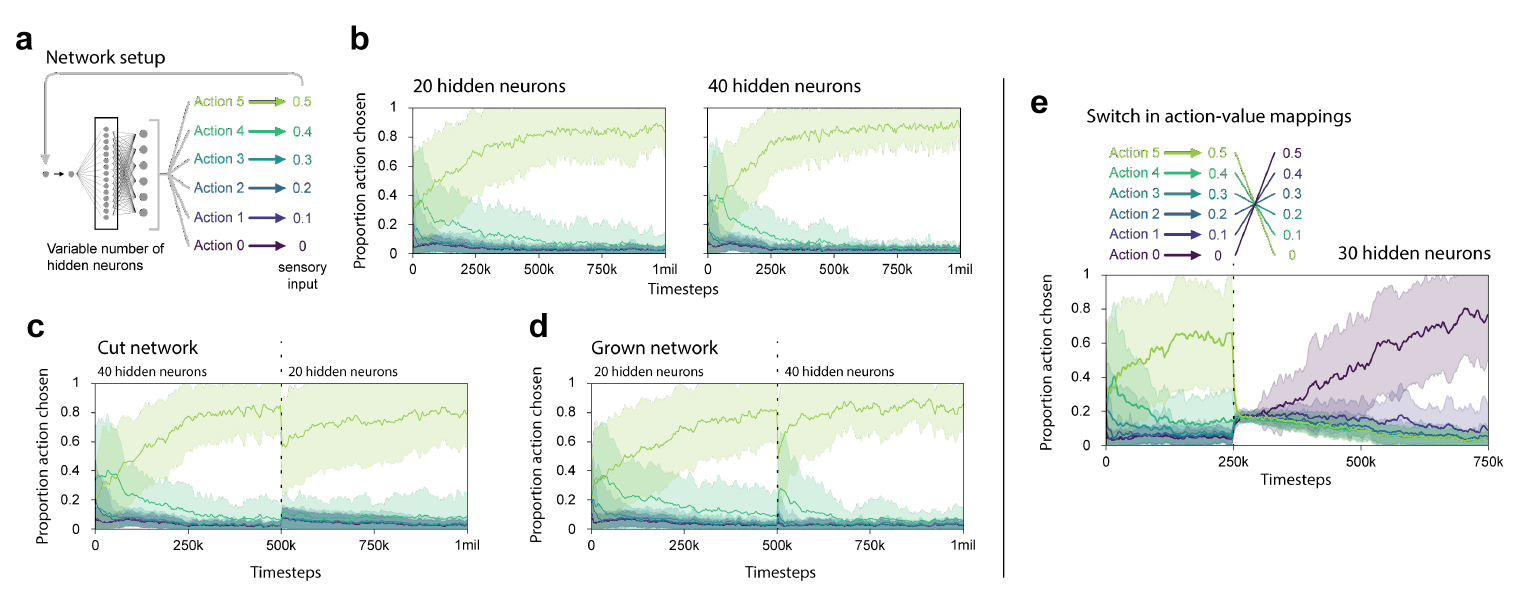
**a**. PaN networks are put in a 6-action environment with linearly spaced rewards. All plots are for 50 random seeds; means and standard deviations in shaded regions. **b**. Networks with 20 (left) or 40 (right) hidden neurons can learn to select the most rewarding action (Action 5). **c**. If networks begin with 40 hidden neurons and are cut to 20 hidden neurons at 500k timesteps, they continue to learn and adapt to the new architecture without external cues. **d**. Likewise, networks adapt when their 20 hidden neurons are doubled to 40 at 500k timesteps. **e**. A network with 30 hidden neurons can adapt to a bandit task that switches reward values midway through the trial. The reward switch occurs at 250k timesteps and the simulations are for 750k timesteps total.

We also support the writing of PaN as a system of SDEs by showing that different step sizes do not noticeably change parameter distributions. Figures 13e-f show plots with a timestep of 1 (see Section D.2.2) and Figures 13g-h show plots with a timestep of 0.05. Again, distributions change very little between conditions.

### D.5 Future work

The SDE picture of PaN opens some avenues of future exploration. In particular, future work could focus on solving the Fokker-Planck equation for the agent’s equilibrium distribution in parameter space, opening the door to the prediction of long-term behavior with arbitrary architectures in any environment. This would allow us to explore agents’ behavior in different tasks with varying number of hidden neurons, changing network connectivities, or with varying learning rates α or ω.

## E Adaptation in bandit tasks

Figure 14 is a series of plots demonstrating continual adaptation, referenced in the main text, Sections 4.3.1 and 4.3.2.

## Footnotes

1 It was important that we made the substitution *s* → *s*(*x*_1_) at this step rather than in the energy function, because if we had made the substitution in the energy function before taking derivatives, we would have inadvertently enforced that *x*_1_ updates to change *s* so that it is closer to *x*_0_. That would have been a non-local interaction, which is not permitted.

