## Supplementary figures and images for "Neuron-level Prediction and Noise can Implement Flexible Reward-Seeking Behavior"

### Video 2

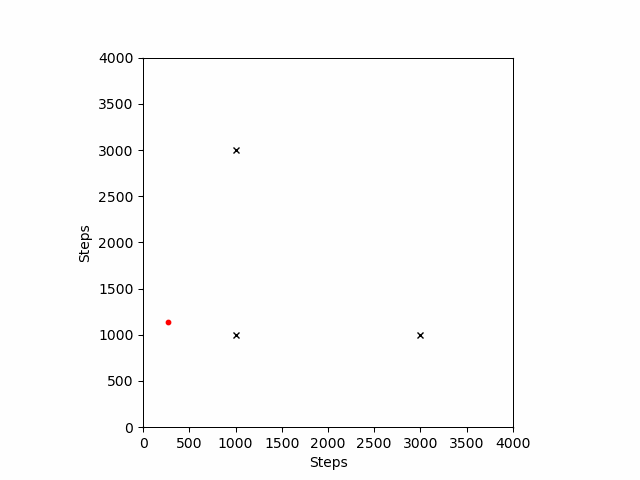
